# Combined *in silico* docking and in vitro antiviral testing for drug repurposing identified lurasidone and elbasvir as SARS-CoV-2 and HCoV-OC43 inhibitors

**DOI:** 10.1101/2020.11.12.379958

**Authors:** Mario Milani, Manuela Donalisio, Rafaela Milan Bonotto, Edoardo Schneider, Irene Arduino, Francesco Boni, David Lembo, Alessandro Marcello, Eloise Mastrangelo

## Abstract

The current emergency of the novel coronavirus SARS-CoV-2 urged the need for broad-spectrum antiviral drugs as the first line of treatment. Coronaviruses are a large family of viruses that already challenged humanity in at least two other previous outbreaks and are likely to be a constant threat for the future. In this work we developed a pipeline based on *in silico* docking of known drugs on SARS-CoV RNA-dependent RNA polymerase combined with in vitro antiviral assays on both SARS-CoV-2 and the common cold human coronavirus HCoV-OC43. Results showed that certain drugs displayed activity for both viruses at a similar inhibitory concentration, while others were specific. In particular, the antipsychotic drug lurasidone and the antiviral drug elbasvir showed promising activity in the low micromolar range against both viruses with good selective index.

## Introduction

The growth in human and animal population density through urbanization and agricultural development, combined with increased mobility and commercial transportation, land perturbation and climate change, all have an impact on virus emergence and epidemiology. Over the past decades, emerging zoonotic RNA viruses continuously gripped the world’s attention, either briefly (like the severe acute respiratory syndrome coronavirus SARS-CoV-1 in 2003), or continuously. Many RNA virus threats were considered as re-emerging including Dengue, Zika, Ebola, and Chikungunya virus, and current consensus predicts that novel and potentially highly pathogenic agents will continue to emerge from the large, genetically variable natural pools present in the environment. Coronaviruses (CoVs) are of particular concern due to high case-fatality rates, lack of therapeutics as well as the ability to seed outbreaks that rapidly cross geographic borders. A large number of highly diverse CoVs have been identified in animal hosts and especially in bat species, where they may have the potential to diffuse in other species including humans (Fan et al., 2019).

Coronaviruses consist of a large and diverse family of viruses that cause multiple respiratory, gastrointestinal and neurologic diseases of varying severity, including the common cold, bronchiolitis, and pneumonia (Weiss & Leibowitz, 2011). The CoV family is divided into four genera (alpha, beta, gamma, and delta) and thus far human CoV are limited to the alpha (HCoV-229E and HCoV-NL63) and beta genera (HCoV-OC43, HCoV-HKU1); the latter includes SARS-CoV-1 and the Middle East respiratory syndrome coronavirus (MERS-CoV). A new previously unknown coronavirus, named SARS-CoV-2, was discovered in December 2019 in Wuhan (Hubei province of China) and sequenced by January 2020 (Lu et al., 2020). SARS-CoV-2 is associated with an ongoing outbreak of atypical pneumonia (COVID-19), and was declared as ‘Public Health Emergency of International Concern’ on January 30^th^, 2020 by the World Health Organization (www.who.int).

Currently, for the COVID19 outbreak, many known drugs are under clinical investigation (Kupferschmidt, 2020) (Magro, 2020), following different general principles and mechanisms of action: 1. the control of cytokine storms due to the hyper-reaction of the immune system against the virus (e.g. corticosteroids, (Salton et al., 2020)); 2. the control of coagulopathy (e.g. heparin) 3. the inhibition of viral RdRp (e.g. prodrugs favipiravir and remdesivir); 4. the inhibition of viral entry (e.g. hydroxychloroquine); 5. the inhibition of the viral main protease (e.g. lopinavir and ritonavir); 5. the inhibition of viral attachment (the viral receptor ACE2 antagonist losartan).

Despite their species diversity, CoVs share key genomic elements that are essential for viral replication, suggesting the possibility to design broad spectrum therapeutic agents to address the current epidemic and manage possible future outbreaks. The target considered in this work to identify new inhibitors is the highly conserved RNA dependent RNA polymerase (RdRp), that plays a crucial role in CoV replication cycle, catalyzing the synthesis of new viral RNA (Te Velthuis et al., 2012). The cryo-EM structure of RdRp of SARS-CoV-1 and of SARS-CoV-2, bound to nsp7 and nsp8 co-factors, have been recently solved (PDB codes: 6NUR (Kirchdoerfer & Ward, 2019), and 6M71 (Gao et al., 2020), respectively). The two proteins share a sequence identity of 96% (98% conservative substitution) and a structural r.m.s.d. of 0.54 Å (considering 788 Cαs).

The exploration of libraries of molecules already in use as human drugs and well characterized in terms of human metabolism might allow the identification of antivirals that could be, in principle, rapidly tested in patients. Accordingly, we chose to analyze *in silico* the public database of approved drugs (DrugBank library, https://go.drugbank.com/), targeting a wide region around the active site of SARS-CoV RdRp. The computational work allowed the selection of 13 commercially available compounds with predicted high affinity for the protein and favorable solubility properties. These potential inhibitors (together with suramin, known to inhibit several RNA viruses) have been tested in cell based assays against SARS-CoV-2 and CoV-OC43 (Su et al., 2016), revealing moderate to high antiviral activities for seven of them. Our results confirm antiviral properties already described for some of the selected compounds, and, more importantly, show new interesting properties for the compounds lurasidone and elbasvir as betacoronavirus inhibitors.

## Results

### *In silico* docking of approved drugs

For the purpose of known drugs repurposing, a total of 6996 molecules were downloaded from the DrugBank library (https://www.drugbank.ca/) to target a wide region (~13,300 Å^3^) around the active site of SARS-CoV RdRp (PDB-ID 6NUR; (Kirchdoerfer & Ward, 2019)). The *in silico* screening was divided into two runs: a fast procedure for the selection of the best 2% of the library (118 compounds), with predicted binding free energy values (ΔG) from −8.9 to −7.6 kcal/mol, followed by a more accurate analysis with AutoDock4.2. In this way we ranked 118 known drugs based on the ΔG value of the best pose for every compound (between −11.7 kcal/mol (predicted Ki=2.7 nM) and −0.12 kcal/mol (predicted Ki=819 mM). The list of the first best 60 compounds is reported in supplemental material (Table S1).

From our list a reasonable number of compounds was selected for cell-based assays (Table 1), taking into account commercial availability and solubility properties. Suramin was added to the list, since it was already known to inhibit several RNA viruses such as flavivirus (Basavannacharya & Vasudevan, 2014) (Albulescu et al., 2017), norovirus (Mastrangelo et al., 2012), but also chikungunya and Ebola viruses (Albulescu et al., 2017) (Henß et al., 2016).

**Table 1.** Compounds selected from *in silico* docking to be experimentally tested

| in silico ranking | Drugbank code | name | predicted ki [nM] for RdRp | 'good' conform. | predicted ki [nM] for main protease |
| --- | --- | --- | --- | --- | --- |
| 5 | DB11363 | Alectinib | 8.5 | 6 | 35.0 |
| 7 | DB09042 | Tedizolid | 20.2 | 4 | 907.6 |
| 9 | DB08901 | Ponatinib | 29.4 | 2 | 32.6 |
| 12 | DB02329 | Carbenoxolone | 34.8 | 17 | 270.4 |
| 13 | DB08815 | Lurasidone | 64.3 | 20 | 7.6 |
| 18 | DB01329 | Cefoperazone | 115.7 | 4 | 71.1 |
| 21 | DB11574 | Elbasvir | 167.6 | 1 | 1.2 |
| 23 | DB00444 | Teniposide | 204.3 | 3 | 68.7 |
| 24 | DB11581 | Venetoclax | 208.4 | 1 | 0.7 |
| 27 | DB00762 | Irinotecan | 247.6 | 2 | 6.7 |
| 32 | DB06290 | Simeprevir | 292.1 | 4 | 31.9 |
| 47 | DB11575 | Grazoprevir | 639.9 | 10 | 40.9 |
| 59 | DB00826 | Natamycin | 945.3 | 16 | 550.9 |

\*number of poses clustered around the one with lower $\Delta G$ (out of 80 poses for each compound).

The best in silico docking pose of each of the selected thirteen compounds, in the RdRp active site, is reported in Figure 1A. The protein region explored is located between thumb, fingers and palm domains and would host growing dsRNA during polymerase activity. Such region defines a wide, complex and variable hydrophilic protein surface, and it is therefore able to host very different types of ligands. In Figure 1B, C, D we report the lurasidone and elbasvir best docking sites, between the thumb and fingers domains and in palm domain, respectively.

**Figure 1.**
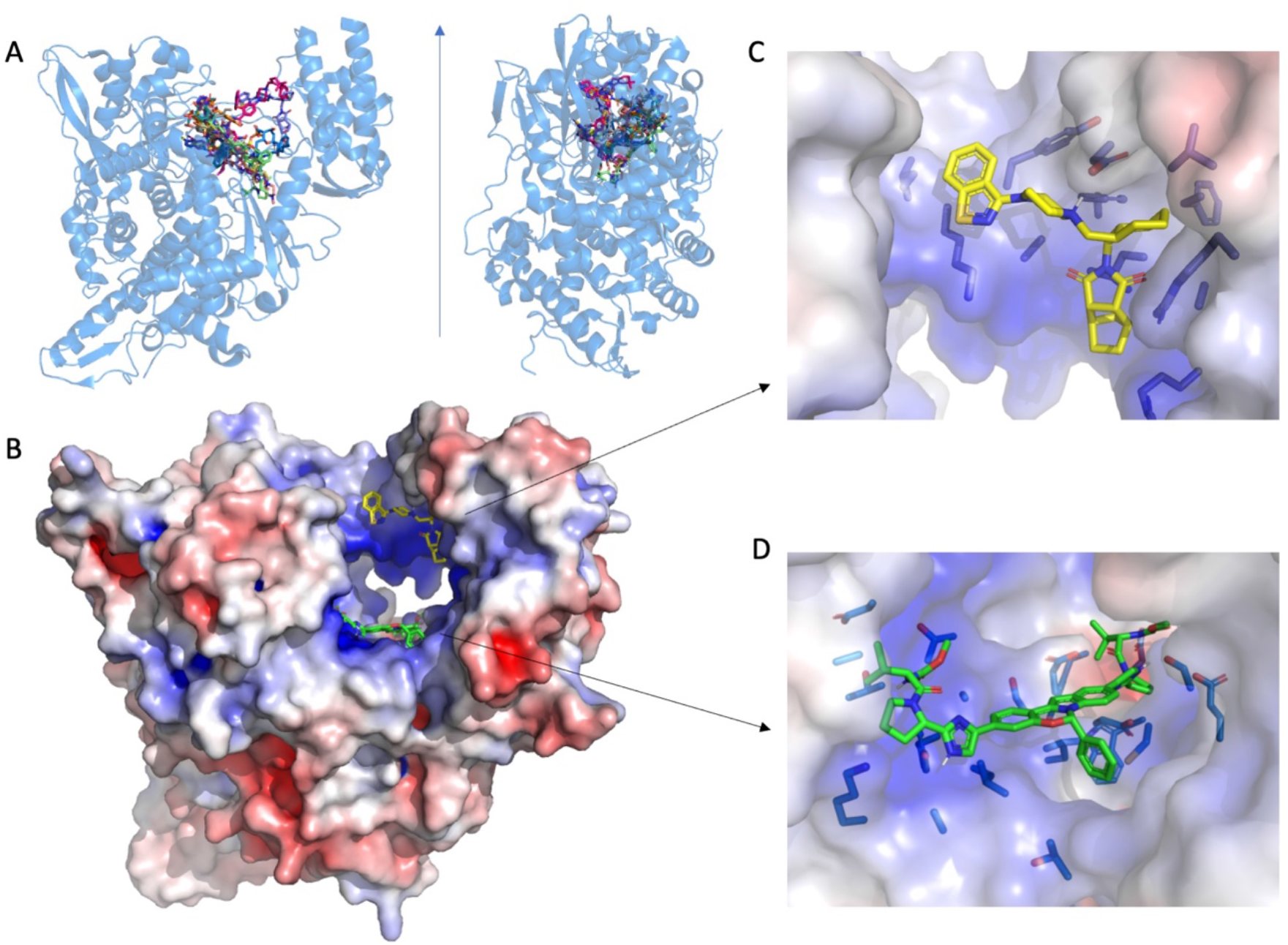
**A)** Best *in silico* docking pose of each of the thirteen selected compounds (sticks with carbon atoms in different colors) in the RdRp (blue transparent cartoons) region around the protein active site that would host growing dsRNA in two perpendicular orientations. **B)** Lurasidone and elbasvir docking site. Lurasidone/elbasvir carbon atoms are shown as yellow/green sticks. The protein surface is colored by electrostatic potential. On **C** and **D** panels, a closer view of the interaction between the 2 compounds and SARS-CoV RdRp, with the protein residues closer to the binding site reported as blue sticks. The figure was prepared using PyMOL (The PyMOL Molecular Graphics System, Version 2.0 Schrödinger, LLC).

Since, among the selected compounds, known inhibitors of viral proteases (like simeprevir and grazoprevir) were also present, we performed an additional *in silico* analysis targeting the active site of main protease (PDB-ID: 6LU7 (Jin et al., 2020)) obtaining the results listed in Table 1. Eight of the selected compounds showed predicted binding affinity for the protease lower than 50 nM suggesting that such known drugs could be in principle active against multiple targets.

### Antiviral activity against SARS-CoV-2 and HCoV-OC43

The antiviral activity of the selected compounds was assessed against two pathogenic CoVs strains: SARS-CoV-2 and HCoV-OC43.

#### Antiviral activity against SARS-CoV-2

A High Content Assay (HCA) has been developed to test antiviral drugs against SARS-CoV-2 *in vitro*. The assay was established using Huh7 cells engineered with the human ACE-2 receptor (Huh7-hACE2) to promote viral infection. The readout of the assay was accessed using immunofluorescence to quantify the number of Spike-positive cells to measure infected cells, and number of nuclei to measure cell viability, as shown in Figure 2.

**Figure 2.**
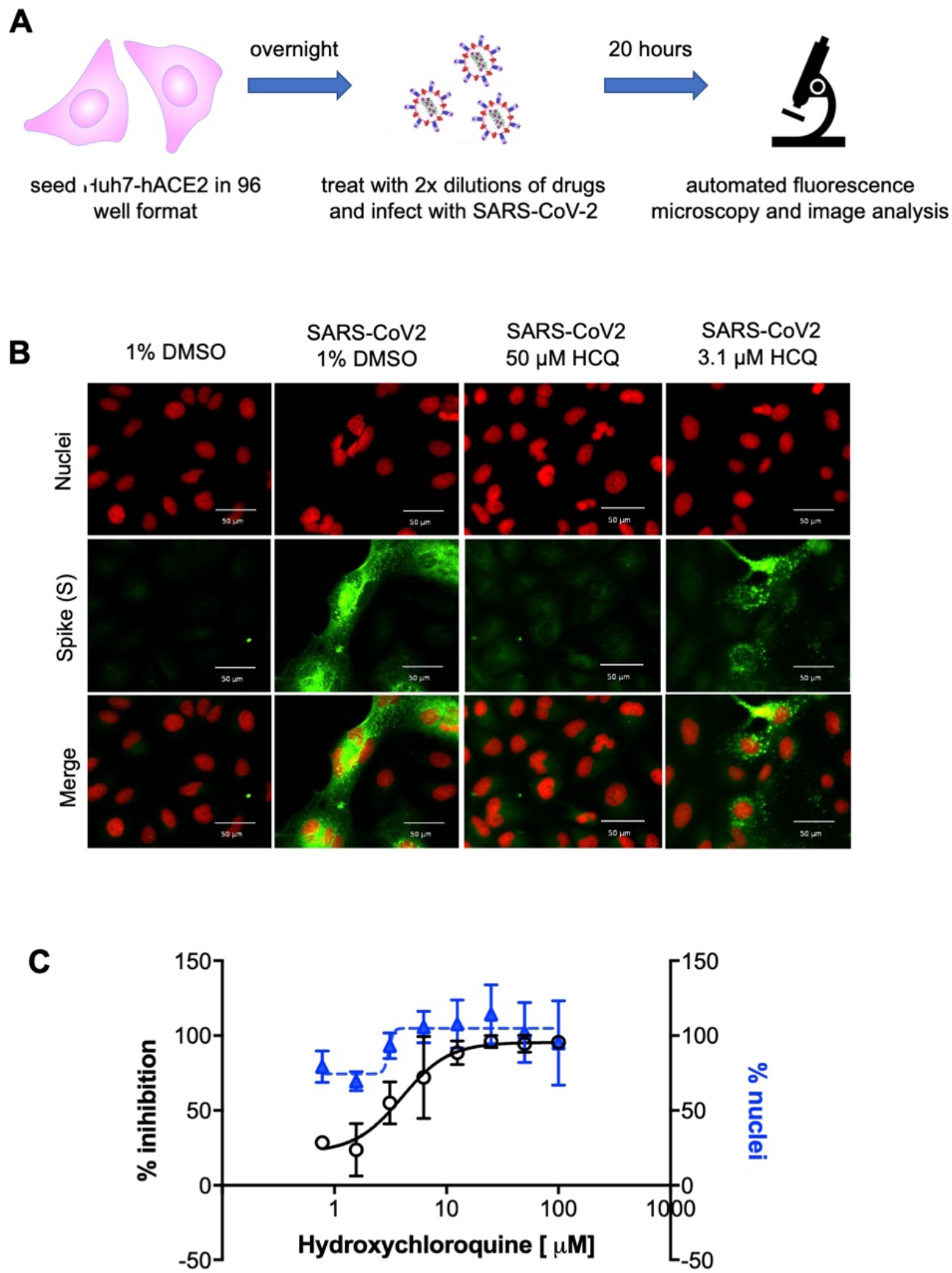
High content screening assay for SARS-CoV-2. **A)** Scheme of SARS-CoV-2 HCS. Huh7-hACE2 were seeded onto 96-well plates, after 24 hours cells were treated with the drugs in two-fold dilutions and immediately infected with SARS-CoV2 (MOI 0.1). 20 hours after infection, cells were fixed, stained and analyzed. **B**) Representative images of the HCS assay with the control drug hydroxychloroquine (HCQ). Nuclei are stained by DAPI (red) and Spike (S) is stained with the mSPI-3022 antibody (green), scale bar corresponds to 50 μm. **C)** Dose response of the positive control hydroxychloroquine. White dots represent the percentage of normalized % of inhibition. Blue triangles represent the % of nuclei compared to the average % of non-infected cells. Error bars represent the standard deviation (SD) of 2 independent experiments.

The panel of compounds was tested in dose response, and the assay was validated using Hydroxychloroquine as reference compound. Results are reported in Table 2, Figure 3 and Figure S2.

**Figure 3.**
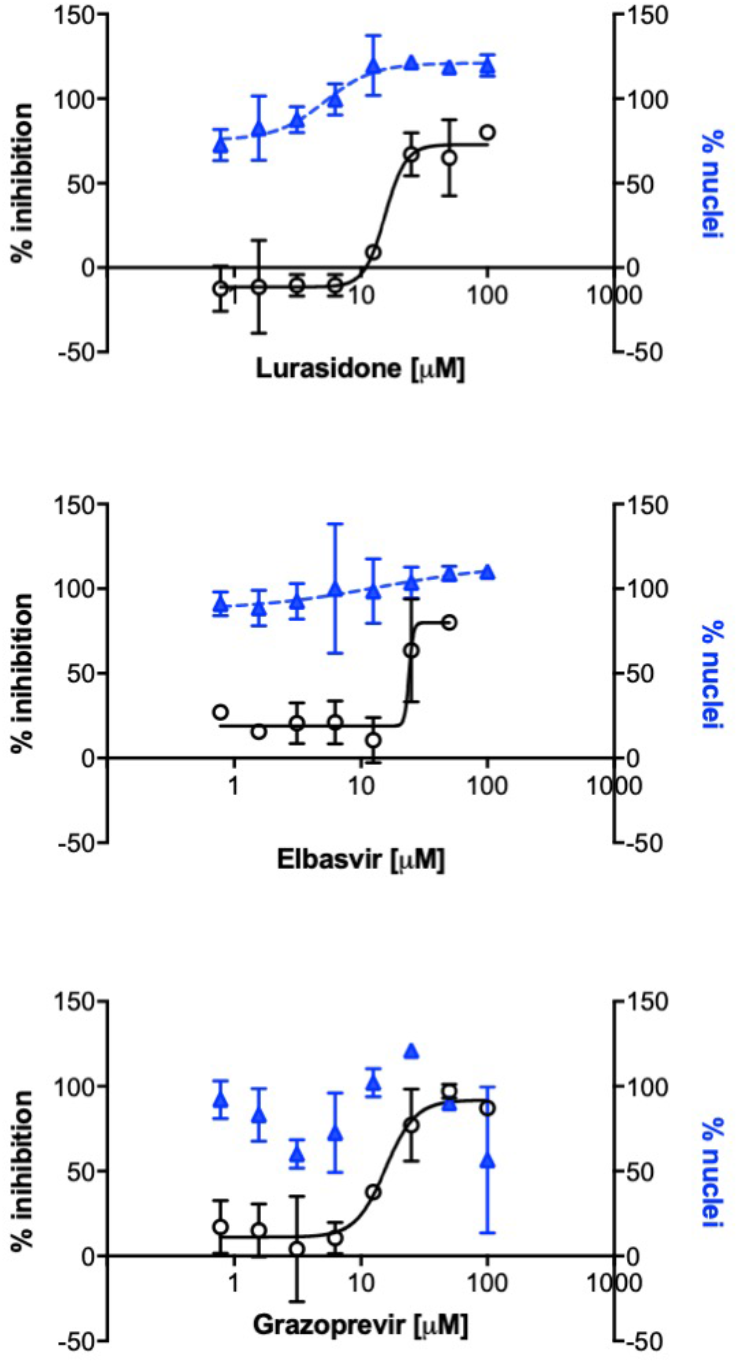
Antiviral efficacy of the best selected compounds against SARS-CoV-2. The antiviral activity was evaluated by the high-content assay infecting Huh7-hACE2 cells exposed to increasing amounts of compounds. Number of nuclei were quantified in parallel. The percentage infectivity inhibition (white dots) was normalized with the average infection ratio of wells treated with 1% DMSO. Percentage of nuclei (blue triangles) was calculated by comparing the average number of nuclei of noninfected wells treated with 1% DMSO. Error bars represent the standard deviation (SD) of 2 independent experiments.

**Table 2.** HCA for the *in silico* selected compounds against SARS-CoV-2

| Compound | EC<sub>50</sub><sup>a</sup> (μM)<br>(mean± SD)<sup>b</sup> | CC<sub>50</sub><sup>c</sup> (μM)<br>(mean± SD) |
| --- | --- | --- |
| Alectinib | *n.a.* | *n.a.* |
| Tedizolid | *n.a.* | *n.a.* |
| Ponatinib | 1.1 ± 0.2 | 8.7 ± 3.8 |
| Carbenoxolone | 66 ± 11 | >100 |
| **Lurasidone** | 18.0 ± 4.6 | >100 |
| Cefoperazone | *n.a.* | *n.a.* |
| **Elbasvir** | 23.0 ± 3.6 | >100 |
| Teniposide | 97.0 ± 0.7 | >100 |
| Venetoclax | 6.2 ± 0.6 | 22.0 ± 6.2 |
| Irinotecan | 85.2 ± 17.0 | >100 |
| Simeprevir | 9.3 ± 2.0 | 47.5 ± 41.0 |
| **Grazoprevir** | 16.0 ± 5.7 | >100 |
| Natamycin | 24.3 ± 4.4 | 35.9 ± 13.4 |
| Suramin | 64.0 ± 6.6 | >100 |

*n.a.* not assessable. <sup>a</sup> Half maximal effective concentration. <sup>b</sup> Standard deviation. <sup>c</sup> Half maximal cytotoxic concentration.

All 14 compounds were tested from a starting concentration of 100 μM in 2-fold dilutions. 11 compounds showed activity at least in one tested concentration. Lurasidone, grazoprevir, venetoclax and elbasvir showed the best outcomes, with EC_50_ in the micromolar range and cytotoxicity >100 μM. Lurasidone showed an EC_50_ of 18 μM with a good dose-response curve devoid of cytotoxicity up to 100 μM. The compound ponatinib reached the lowest EC_50_ (1.1 μM) against SARS-CoV2, although the elevated cytotoxicity (CC_50_=8.7 μM) indicated a poor selectivity index (calculated as the ratio of the CC_50_ and the EC_50_ values).

The known HCV inhibitors elbasvir, simeprevir and grazoprevir showed EC_50_ values of 23 μM, 9.3 μM and 16 μM, respectively. However, their activity as HCV inhibitors is in the low nanomolar range and simeprevir showed an unfavorable CC_50_ of 47.5 μM. The antiviral activity of suramin was confirmed, with an EC_50_ of 64 μM, similarly to previous reports (Salgado-Benvindo et al., 2020). A weak activity was detected with compounds irinotecan, teniposide and carbenoxolone, all with an EC_50_ around 50-100 μM. In conclusion, our data showed *in vitro* activity for most of the compounds selected *in silico* against SARS-CoV RdRp. Among all, lurasidone grazoprevir and elbasvir showed the best antiviral profile against SARS-CoV-2 (Figure 3).

#### Antiviral activity against HCoV-OC43

In order to evaluate the anti-HCoV-OC43 activity of the selected compounds, focus reduction assays were performed on MRC-5 cells, as described in the Materials and Methods section and elsewhere (Marcello et al., 2020). Results on antiviral activity and cell toxicity are reported in Table 3, Figure 4 and Figure S3.

**Figure 4.**
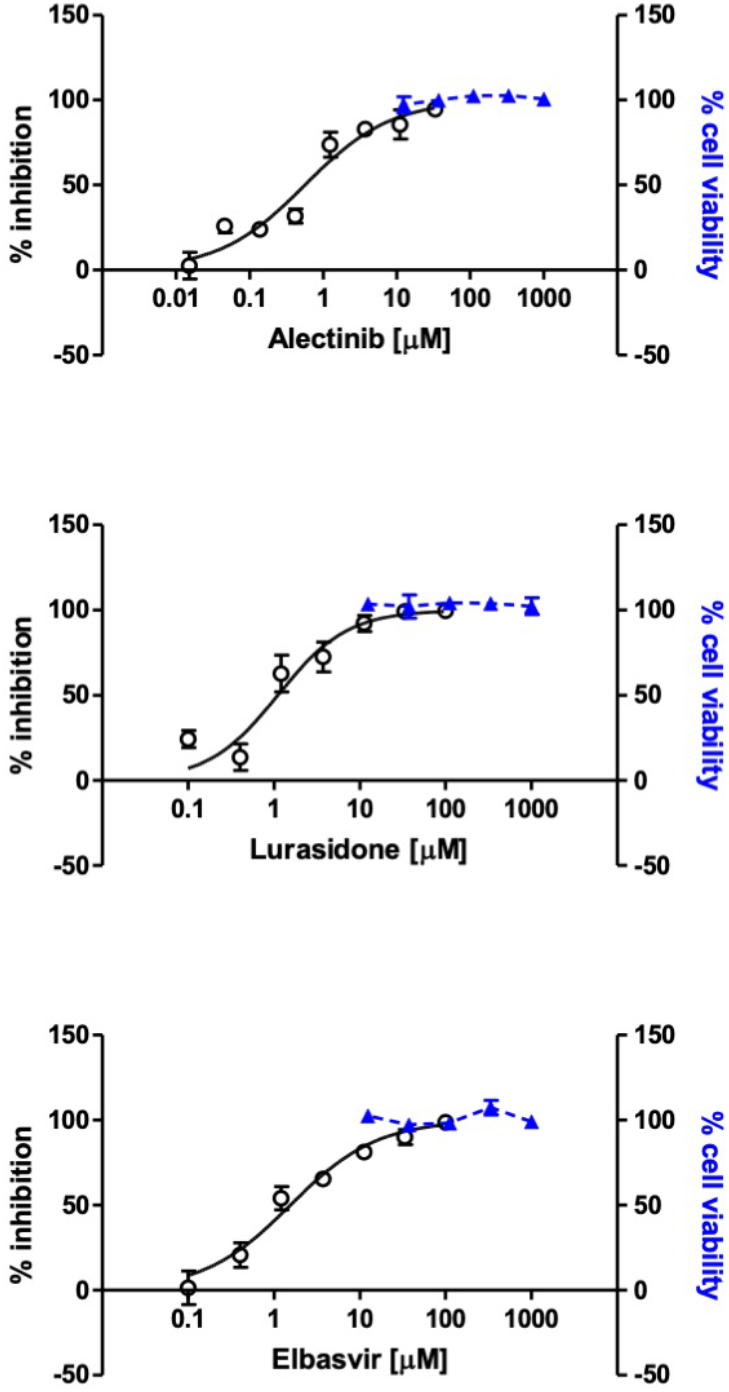
Antiviral efficacy of the selected compounds against HCoV-OC43. The antiviral activity of compounds was evaluated by focus reduction assay, infecting MRC-5 cells in presence of increasing concentration of compounds. Cell viability assays were performed in the same conditions as for antiviral assays, in absence of viral inoculum. The percentage infectivity inhibition (white dots) and the percentage of cell viability (blue triangles) were calculated by comparing treated and untreated wells. Error bars represent the standard deviation (SD) of 3 independent experiments.

**Table 3:** Anti-HCoV-OC43 activity of the selected compounds.

| Compound | EC<sub>50</sub><sup>a</sup> (μM)<br>(mean± SD)<sup>b</sup> | CC<sub>50</sub><sup>c</sup> (μM)<br>(mean± SD) |
| --- | --- | --- |
| **Alectinib** | 0.6 ± 0.2 | >1000 |
| Tedizolid | 94.0 ± 25 | >1000 |
| **Ponatinib** | 0.10 ± 0.05 | 3.1 ± 0.5 |
| Carbenoxolone | 45.7 ± 8.0 | 251 ± 17 |
| **Lurasidone** | 1.1 ± 0.4 | >1000 |
| Cefoperazone | n.a. | >1000 |
| **Elbasvir** | 1.5 ± 0.5 | >1000 |
| Teniposide | n.a. | 701 ± 281 |
| Venetoclax | 3.5 ± 0.5 | 10.6 ± 1.5 |
| Irinotecan | n.a. | 63.4 ± 11.4 |
| Simeprevir | 2.1 ± 0.6 | 11.1 ± 2.3 |
| Grazoprevir | 11.0 ± 1.1 | 70.0 ± 12.8 |
| Natamycin | n.a. | 47.3 ± 4.4 |
| Suramin | 19.3 ± 4.3 | >1000 |

n.a. not assessable. <sup>a</sup> Half maximal effective concentration. <sup>b</sup> Standard deviation. <sup>c</sup> Half maximal cytotoxic concentration.

Among the tested compounds, alectinib showed the strongest inhibitory activity against HCoV-OC43, with an EC_50_ in the low micromolar range (0.6 μM). Lurasidone and elbasvir also exerted high antiviral activity against HCoV-OC43, exhibiting EC_50_ in the low micromolar range: 1.1 μM and 1.5 μM, respectively. A moderate antiviral activity was shown by tedizolid, carbenoxolone and suramin, with EC_50_ ranging from 11.0 μM to 94.0 μM. The afore-mentioned compounds’ antiviral effect was not a consequence of cytotoxicity, since none of the screened compounds significantly reduced cell viability at any concentration used in the antiviral assays (i.e. up to 100 μM), exhibiting CC_50_ values higher than 1000 μM. By contrast, the remaining compounds did not exhibit interesting features as anti-HCoV-OC43 molecules, due to either no antiviral activity (cefoperazone, teniposide, irinotecan, natamycin), or low-moderate selectivity index (ponatinib, venetoclax, simeprevir, grazoprevir). In summary, these data showed that alectinib, lurasidone and elbasvir were endowed with strong anti-HCoV-OC43 activity (Figure 4), with minimal toxicity and selectivity indexes higher than 600.

## Discussion

An accurate *in silico* docking search within a wide region around the SARS-CoV RdRp active site, allowed us to select 13 known drugs from the DrugBank library to be experimentally tested. We added suramin to the list, a well-known compound able to inhibit different RNA viruses (Mastrangelo et al., 2012) (De Clercq, 1979) (Albulescu et al., 2015).

Unexpectedly, a rather high percentage (>60%) of the selected compounds showed some *in vitro* activity against one or both of the tested CoV strains. A possible explanation for such a positive result is related to the characteristics of the protein region selected for the *in silico* docking. Such a portion of the protein is a wide, complex and rather hydrophilic surface with many conformational degrees of freedom, allowing it to adapt to the growing dsRNA during translation. Accordingly, the average crystallographic (or cryoEM) conformation of this region must be capable to accommodate different kind of ligands (as shown in Figure 1), with a preferential affinity for large compounds possessing polar/charged moieties and planar aromatic groups: i.e. compounds that generally mimics RNA backbone and bases. In other words, the *in silico* docking on RdRp not only selects compounds potentially capable of interfering with the polymerase activity but could also act as a molecular filter for the selection of properties generally favorable for protein binding/inhibition. This explanation is supported by the predicted high affinity for the main protease of most of the selected compounds (Table 1).

Among the tested compounds lurasidone and elbasvir displayed higher activity and lower cytotoxicity against both SARS-CoV-2 and HCoV-OC43 strains. Lurasidone lead to complete inhibition of both strains with EC_50_ values in the micromolar range (18 and 1.1 μM, respectively) and favorable selectivity indexes. Lurasidone is an antipsychotic drug for treatment of acute depression and schizophrenia, known to bind with a low nanomolar affinity to Dopamine-2, 5-HT1A, 5-HT2A, and 5-HT7 receptors, and with slightly lower affinity to alpha-2C adrenergic receptors (Greenberg & Citrome, 2017). Lurasidone was already identified as a potential inhibitor of SARS-CoV-2 main protease (Elmezayen et al., 2020) and in our in silico analysis it showed good predicted binding affinity for both RdRp and main protease. Since SARS-CoV-2 and HCoV-OC43 share a high level of protein sequence conservation (Vijgen et al., 2005) we hypothesize that mechanisms of action of lurasidone against these viruses might be the same.

Elbasvir inhibited SARS-CoV-2 and HCoV-OC43 with EC_50_ values in the micromolar range (about 23 and 1.5 μM, respectively). Previous *in silico* studies predicted elbasvir as a high affinity compound for the RdRp, the papain-like protease and the helicase of SARS-CoV-2 (Balasubramaniam & Shmookler Reis, 2020), whereas our *in silico* investigation suggested a preferential binding for the main protease (1.2 nM). Elbasvir is an inhibitor of the HCV NS5A protein that has not homologues in coronaviruses: in light of our and previous work results, it has the potential to inhibit different viral proteins.

Alectinib (with the best *in silico* Ki against RdRp) showed no activity against SARS-CoV-2 but it is the best compound against HCoV-OC43 (EC_50_=0.6 μM, CC_50_ value >1000 μM). Alectinib inhibits the anaplastic lymphoma kinase (ALK) tyrosine kinase receptor in the nM range (Kinoshita et al., 2012), binding to the ATP binding site. Therefore, we can speculate that it might inhibit other kinases essential for HCoV-OC43 replication in MRC-5 cells. Moreover, we can speculate that the compound could act as a competitive inhibitor of RdRp, interfering with the nucleotide binding site(s) of the enzyme. In this case a possible explanation for the different results obtained for SARS-CoV-2 and HCoV-OC43 could be related to the differences between the two polymerases (sequence identity of 55.1%) (Elfiky, 2020).

Grazoprevir inhibited SARS-CoV-2 with an EC_50_ around 16 μM (CC_50_ value >100 μM). Published computational studies suggested grazoprevir as a potential inhibitor of the nucleocapsid protein or the papain-like protease of SARS-CoV-2 (Behera et al., 2020). Grazoprevir is an inhibitor of HCV protease and it is often used for therapy in combination with elbasvir (in the drug named zepatier).

Simeprevir, another inhibitor of HCV protease, has been previously shown to inhibit SARS-CoV-2 in synergy with remdesivir (Lo et al., 2020). In our experiments it showed a similar potency against both SARS-CoV-2 (EC_50_ about 9.3 μM) and HCoV-OC43 (EC_50_ about 2.1 μM) but with low SI. From our docking results its effect is likely directed against the protease, but RdRp inhibition cannot be excluded.

ABT-199, also known as venetoclax, is a potent selective Bcl2 inhibitor, which induces the apoptosis pathway. An early work showed that Bcl2 expression prevents SARS-CoV induced apoptosis (Bordi et al., 2006). In addition, previous reports demonstrated that SARS-CoV 7a protein was dependent on Bcl2 to induce apoptosis, suggesting Bcl2 as an important host factor for virus replication and pathogenesis (Tan et al., 2007). However, despite its good EC_50_ (about 6.2 μM) against SARS-CoV-2, venetoclax shows high toxicity in the tested cells.

Ponatinib, an oral drug for the treatment of chronic myeloid leukemia and Philadelphia chromosome-positive acute lymphoblastic leukemia, was already proposed as SARS-CoV-2 inhibitor (Nguyen et al., 2020) (Sauvat et al., 2020) (Gordon et al., 2020), and it is shown here to inhibit SARS-CoV-2 and HCoV-OC43 with a poor selective index.

All the other compounds, although some of them have been described in the literature as potential inhibitors of SARS-CoV-2 (i.e. teniposide (Kadioglu1 et al., n.d.) and irinotecan (B, 2020), did not show any relevant activity in either of the two viruses tested.

### Conclusions

In our work we have: 1. excluded SARS-CoV-2 antiviral activity for teniposide (Kadioglu1 et al., n.d.) and irinotecan (B, 2020), selected from previous computational studies; 2. showed the ability of some of the already described anti-SARS-CoV-2 compounds to inhibit also coronavirus HCoV-OC43 causing the common cold (suramin, ponatinib - although with a low SI); and most importantly 3. showed the capability of some of the selected drugs to selectively inhibit HCoV-OC43 (alectinib) or SARS-CoV-2 (grazoprevir) or be active against both CoV strains (lurasidone and elbasvir). Treatment of CoV infections with drugs that could inhibit different viral targets, as predicted for lurasidone and elbasvir, would be an effective way to lower chances of the emergence of drug resistant viral strains.

Of note, in previous works it was demonstrated that alectinib (Song et al., 2015) could penetrate the blood-brain barrier (BBB) exerting its activity in the central nervous system (CNS). Since HCoV-OC43, as other coronaviruses, is able to invade the CNS (Dubé et al., 2018), alectinib might be an interesting candidate for the treatment of HCoV-OC43 persistent infections in the brain. Moreover, the free levels of alectinib found in both plasma and cerebrospinal fluid are similar (Herden & Waller, 2018) and its EC_50_ against HCoV-OC43 (0.6 uM) is lower than the maximum level attainable in human serum with daily recommended dosage (676 ng/mL corresponding to 1.4 μM) (Ly et al., 2018).

In conclusion, our approach allowed the identification of lead-drugs for further *in vitro* and clinical investigation to contain the present outbreak. Furthermore, it could contribute to the identification of broad spectrum anti-CoV inhibitors / therapies that would allow for a rapid and effective reaction to future epidemics.

## Materials and Methods

### In silico docking

The virtual Library of DrugBank (https://www.drugbank.ca/) employed for the docking analysis (6996 compounds) includes commercially available FDA-approved drugs as well as experimental drugs going through the FDA approval process. The atomic coordinates of SARS-CoV RdRp (PDB-ID 6NUR) bound to NSP7 and NSP8 cofactors, were chosen as docking model for CoV polymerase. Hydrogen atoms and Kollman charges (Singh & Kollman, 1984) were added using the program Python Molecule Viewer 1.5.4 (MGL-tools package http://mgltools.scripps.edu/). The protein model was then used to build a discrete grid within a box of dimensions 22.5×26.3×22.5 Å^3^ (program autogrid (Goodford, 1985)) as the explored volume for the docking search. The grid was centered near the side chain of Lys545, to include a wide region around the protein active site. During the computational analysis, the protein was constrained as rigid, whereas the small molecules were free to move. The *in silico* screen was divided into two runs: a fast procedure using AutoDock Vina (Trott & Olson, 2009) for the selection of the best compounds, followed by a more accurate screen using AutoDock4.2 (Morris et al., 2009). The Autodock Vina docking search produced a ranked list of all compounds, with predicted binding free energy values (ΔG) ranging between −0.9 kcal/mol and −8.9 kcal/mol. The best 118 compounds (~2% of the library, ΔG between −7.6 to −8.9 kcal/mol) were further analyzed using AutoDock4.2 (Morris et al., 2009), with 80 hybrid GA-LS genetic algorithm runs. Among the molecules with higher predicted affinity for RdRp (ΔG values varying between −0.12 and −11.7 kcal/mol), 13 FDA approved drugs were selected, taking into account commercial availability and solubility properties, for *in vitro* assays. Since among such drugs were present known inhibitor of viral protease we investigated their binding affinity for CoV main protease (PDB-ID 6LU7; (Jin et al., 2020)). Briefly, we explored with AutoDock4.2 a region of 15×22.5×22.5 Å^3^ (after mutating the active site Cys145 to Ala) centered between the side chains of Asn142 and Gln189.

### SARS-CoV-2 cell based assays

#### Cell lines and viruses

Vero E6 cells (ATCC-1586), the human hepatocarcinoma Huh7 cells kindly provided by Ralf Bartenschlager (University of Heidelberg, Germany) and Huh 7 engineering by lentivirus transduction to overexpress the human ACE2 (Huh-7hACE2) were cultured in Dulbecco’s modified 7 Eagle’s medium (DMEM) supplemented with 10% fetal bovine serum (FBS, Gibco). Working stocks of SARS-CoV-2 ICGEB-FVG_5 isolated in Trieste, Italy, were routinely propagated and titrated on Vero E6 cells (Licastro et al., 2020).

#### Compounds preparation

Compounds were prepared in 2-fold serial dilutions (8 points dilutions) in DMSO, and then diluted 16x in PBS in an intermediate plate. Finally, compounds were transferred to the 96 well assay plate containing cells and virus medium (6x in grown medium, final dilution 100x).

#### High Content Assay

Huh 7-hACE2 cells were seeded in a 96 wells’ plate, at 8×10^3^ cells/well density and incubated at 37°C overnight. Cells were treated with serial dilution of the compounds andthen infected with SARS-CoV-2 at 0.1 MOI. Controls included: positive controls like infected cells treated with 50 μM of Hydroxychloroquine as well as non-infected cells treated with vehicle (1 % DMSO), and negative controls such as infected cells treated with vehicle. Plates were incubated for 20 h at 37°C, and then fixed with 4% PFA for 20 min at room temperature and washed twice with PBS 1x. Cells were treated with 0.1% of Triton-X for 15 min, followed incubation of 30 min in blocking buffer (PBS containing 1% of bovine serum albumin-BSA). Then, a primary recombinant monoclonal Spike antibody (CR3022) was diluted in blocking buffer and incubate for 2 h at 37°C (Rajasekharan et al., 2020). Cells were washed 2 times in PBS and incubated the secondary antibody AlexaFluor488-conjugated goat anti-mouse IgG (Cat No. A-11001, Thermo-Scientific) plus DAPI for 1 h at 37°C. Each plate was washed twice with PBS. All plates were filled up with 150 μl of PBS/well. Digital images were acquired using a high content imaging system, the Operetta (Perkin Elmer). The digital images were taken from 9 different fields of each well at 20× magnification. Total number of cells and the number of infected cells were analysis using Columbus Image Data Storage and Analysis System (Perkin Elmer).

#### Data normalization and analysis

Infection ration was defined as ratio between (i) the total number of infected cells, and (ii) the total number of cells. Data were normalized with the negative (DMSO-treated, infected cells) and positive (infected cells treated with 50 μM Hydroxychloroquine) controls. Percentage inhibition was calculated based in infection ratio values with the formula: (1-(infection ratio samples – Avarage (Av) infection ratio of positive control) /(Av. infection ratio of negative control – Av. infection ratio of positive control)) x100. Percentage of nuclei was calculate from values of cell number with the formula: Cell number test sample/Avg. cell number of positive control) × 100. Values were plotted against dilutions expressed as antilog. The half maximal effective concentration (EC_50_) and the half maximum cytotoxic concentration (CC_50_) were calculated using GraphPad Prism Version 7.

### HCoV-OC43 cell-based assays

#### Reagents

Dimethyl sulfoxide (DMSO) was purchased from Sigma-Aldrich (Saint Louis, MO). The mouse anti-coronavirus monoclonal antibody MAB9013 was purchased from Merck (Darmstadt, Germany). The secondary antibody peroxidase-conjugated AffiniPure F (ab’)2 Fragment Goat Anti-Mouse IgG (H+L) was purchased from Jackson ImmunoResearch Laboratories Inc. (West Grove, PA, USA).

#### Cell lines and viruses

Human lung fibroblast cells MRC-5 (ATCC^®^ CCL-171) were propagated in Dulbecco’s Modified Eagle Medium (DMEM; Sigma, St. Louis, MO, USA) supplemented with 1% (v/v) penicillin/streptomycin solution (Euroclone, Milan, Italy) and heat inactivated, 10% (v/v) fetal bovine serum (Sigma). Human coronavirus strain OC43 (HCoV–OC43) (ATCC^®^ VR-1558) was purchased from ATCC (American Type Culture Collection, Rockville, MD, USA). The virus was propagated in MRC-5 cells at 33°C, in a humidified 5% CO_2_ incubator, and titrated by standard plaque method on MRC-5 cells, as described elsewhere (Marcello et al., 2020); titers were expressed in terms of plaque forming units per ml (PFU/ml).

#### Cell viability assay

Cell viability was measured using the MTS assay, as described elsewhere (Lembo et al., 2014). MRC-5 cells were seeded at a density of 2×10^4^ cells/well in 96-well plates and treated the next day with compounds at concentrations ranging from 1000 to 0.05 μM, under the same experimental conditions described for the antiviral assays. Treatment of control wells with equal volumes of DMSO was performed in order to rule out the possibility of any cytotoxic effect ascribable to the solvent. After 20 h of incubation, cell viability was determined using the Cell Titer 96 Proliferation Assay Kit (Promega, Madison, WI, USA) according to the manufacturer’s instructions. Absorbances were measured using a Microplate Reader (Model 680, Bio-Rad Laboratories, Hercules, CA, USA) at 490 nm. The effect on cell viability at different concentrations of compounds was expressed as a percentage, by comparing absorbances of treated cells with those of cells incubated with culture medium and equal volumes of DMSO. The 50% cytotoxic concentrations (CC_50_) and standard deviation (SD) values were determined using GraphPad Prism 5.0 software (GraphPad Software, San Diego, CA).

#### Antiviral assay

The antiviral activity was determined by focus reduction assay. MRC-5 cells were seeded, at 2×10^4^ cells/well density, in 96-well plates and incubated at 37°C overnight. The next day, the medium was removed from the plates and infection was performed with ca. 40 PFU of a stock of HCoV-OC43 (MOI 0.2 PFU/cells) in presence of serial dilutions of compounds, ranging from 100 to 0.005 μM. Control wells were infected in presence of equal volumes of DMSO. After 20 h of incubation at 33°C in a humified 5% CO_2_ atmosphere, cells were fixed with cold acetone-methanol (50:50) and subjected to indirect immunostaining by using an anti-coronavirus monoclonal antibody (MAB9013). The number of immunostained foci was counted, and the percent inhibition of virus infectivity was determined by comparing the number of foci in treated wells with the number in untreated control wells. The focus reduction assays were conducted in three independent experiments. Where possible, half-maximal antiviral effective concentration (EC_50_) and SD values were calculated by regression analysis using the software GraphPad Prism 5.0 (GraphPad Software, San Diego, CA) by fitting a variable slope-sigmoidal dose–response curve.

#### Statistical analyses

All data were analyzed using GraphPad Prism 5.0 (GraphPad Software, San Diego, CA). All results are presented as means ± standard deviations.

## Acknowledgments

We would like to thank all the people that contributed to the #FarmaCovid crowd founding initiative and in particular Mrs. Paola Allegretti, who with great commitment has advised many people from Perugia to support our research work.

