## supplemental data for "Combined *in silico* docking and in vitro antiviral testing for drug repurposing identified lurasidone and elbasvir as SARS-CoV-2 and HCoV-OC43 inhibitors"

Table S1. List of the first best 60 compounds selected by in silico docking. In red the compounds selected for cell-based assays.

| in silico ranking | Cpd | name | predicted ki [nM] |
|---|---|---|---|
| 1 | DB09534 | Ecamsule | 2.7 |
| 2 | DB02633 | Cibacron Blue | 3.1 |
| 3 | DB01830 | AP-22408 | 4.2 |
| 4 | DB01879 | *experimental* | 4.3 |
| **5** | **DB11363** | **Alectinib** | **8.5** |
| 6 | DB04330 | Bilh 434 | 12.2 |
| **7** | **DB09042** | **Tedizolid** | **20.2** |
| 8 | DB03276 | *experimental* | 26.7 |
| **9** | **DB08901** | **Ponatinib** | **29.4** |
| 10 | DB06581 | Bevirimat | 29.6 |
| 11 | DB04016 | *experimental* | 30.0 |
| **12** | **DB02329** | **Carbenoxolone** | **34.8** |
| **13** | **DB08815** | **Lurasidone** | **64.3** |
| 14 | DB08823 | Spinosad | 71.8 |
| 15 | DB05340 | ATL-2502 | 86.1 |
| 16 | DB07189 | *experimental* | 110.2 |
| 17 | DB09335 | Alatrofloxacin | 115.3 |
| **18** | **DB01329** | **Cefoperazone** | **115.7** |
| 19 | DB01396 | Digitoxin | 148.6 |
| 20 | DB03712 | RU85053 | 164.9 |
| **21** | **DB11574** | **Elbasvir** | **167.6** |
| 22 | DB06210 | Eltrombopag | 195.9 |
| **23** | **DB00444** | **Teniposide** | **204.3** |
| **24** | **DB11581** | **Venetoclax** | **208.4** |
| 25 | DB09233 | Cronidipine | 212.8 |
| 26 | DB02432 | RU90395 | 216.7 |
| **27** | **DB00762** | **Irinotecan** | **247.6** |
| 28 | DB01988 | *experimental* | 251.2 |
| 29 | DB02169 | *experimental* | 259.4 |
| 30 | DB04289 | Genz-10850 | 279.2 |

| in silico ranking | Cpd | name | predicted ki [nM] |
|---|---|---|---|
| 31 | DB07213 | *experimental* | 289.6 |
| **32** | **DB06290** | **Simeprevir** | **292.1** |
| 33 | DB01836 | *experimental* | 376.4 |
| 34 | DB05868 | Ciluprevir | 390.4 |
| 35 | DB00511 | Acetyldigitoxin | 416.1 |
| 36 | DB09280 | Lumacaftor | 454.3 |
| 37 | DB01251 | Gliquidone | 464.8 |
| 38 | DB06925 | *experimental* | 485.7 |
| 39 | DB04888 | Bifeprunox | 519.3 |
| 40 | DB03957 | SP2456 | 526.8 |
| 41 | DB04881 | Elacridar | 527.0 |
| 42 | DB08512 | *experimental* | 590.0 |
| 43 | DB02112 | Zk-806450 | 596.1 |
| 44 | DB02194 | *experimental* | 602.5 |
| 45 | DB04698 | *experimental* | 618.0 |
| 46 | DB01267 | Paliperidone | 632.9 |
| **47** | **DB11575** | **Grazoprevir** | **639.9** |
| 48 | DB00872 | Conivaptan | 696.8 |
| 49 | DB04673 | *experimental* | 697.5 |
| 50 | DB04877 | Voacamine | 709.5 |
| 51 | DB07847 | *experimental* | 726.0 |
| 52 | DB01897 | *experimental* | 740.9 |
| 53 | DB03933 | C-1027 | 799.3 |
| 54 | DB00773 | Etoposide | 801.5 |
| 55 | DB08164 | *experimental* | 803.0 |
| 56 | DB08237 | *experimental* | 850.0 |
| 57 | DB08676 | *experimental* | 855.5 |
| 58 | DB00430 | Cefpiramide | 864.2 |
| **59** | **DB00826** | **Natamycin** | **945.3** |
| 60 | DB01459 | Bezitramide | 953.5 |

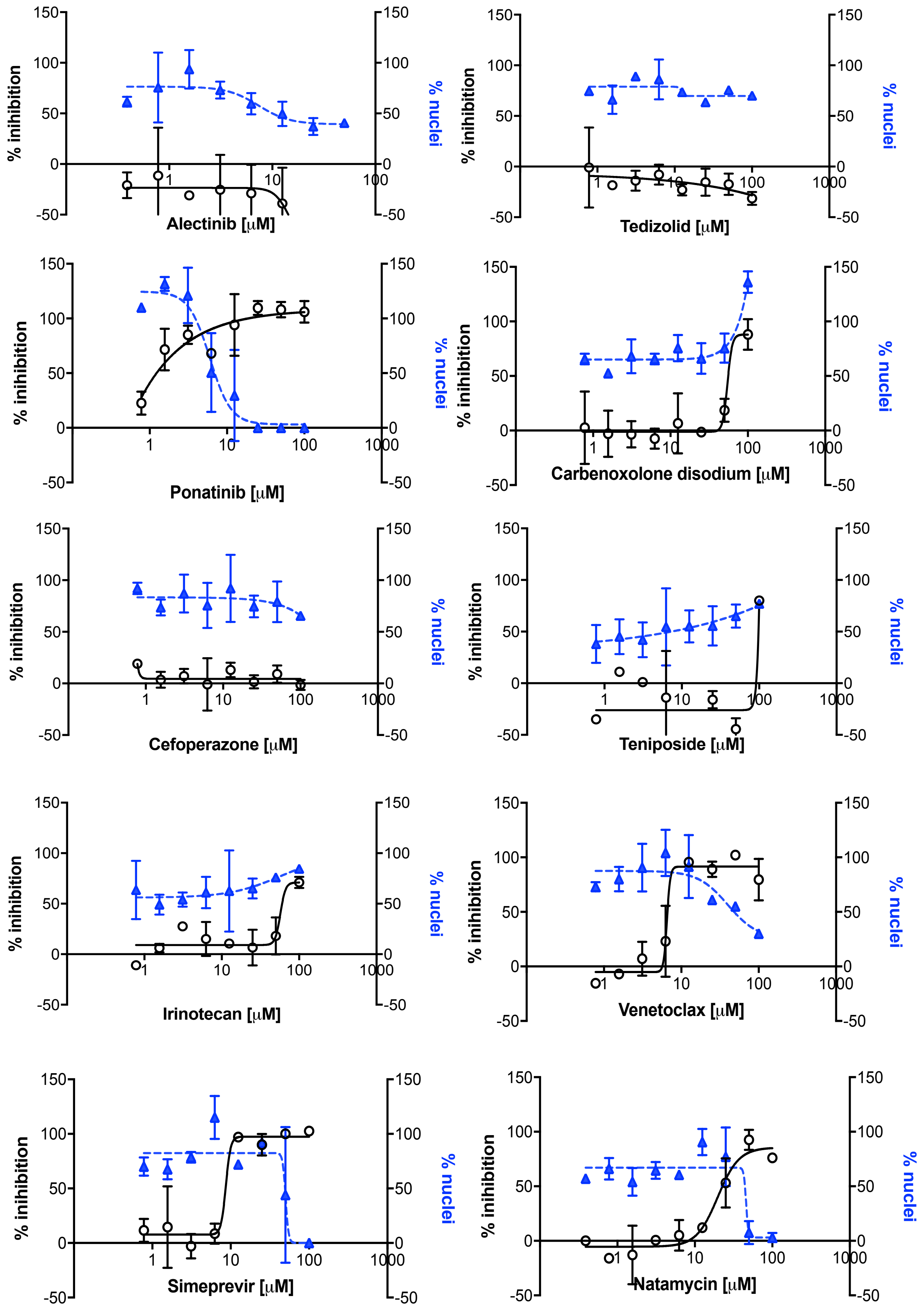

% inihibition
% nuclei
Alectinib [μM]
Tedizolid [μM]
Ponatinib [μM]
Carbenoxolone disodium [μM]
Cefoperazone [μM]
Teniposide [μM]
Irinotecan [μM]
Venetoclax [μM]
Simeprevir [μM]
Natamycin [μM]
150
100
50
0
-50
1
10
100
1000

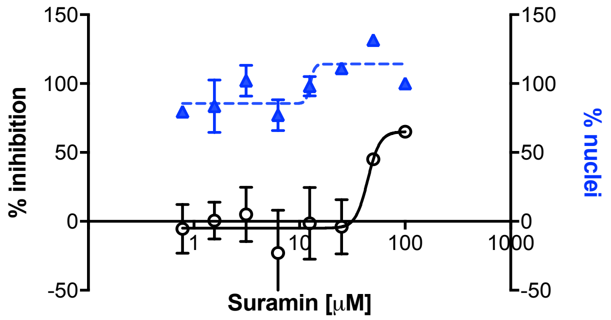


Figure S1: Antiviral efficacy of the selected compounds against SARS-CoV-2. The antiviral activity of compounds was evaluated by means of focus reduction assay, infecting Huh7-hACE2 cells in presence of increasing concentration of compounds. Number of nuclei were quantified in parallel. The percentage infectivity inhibition (white dots) was normalized with the average infection ratio of wells treated with 1% DMSO. Percentage of nuclei (blue triangles) was calculated by comparing the average number of nuclei of noninfected wells treated with 1% DMSO. Error bars represent the standard deviation (SD) of 2 independent experiments.

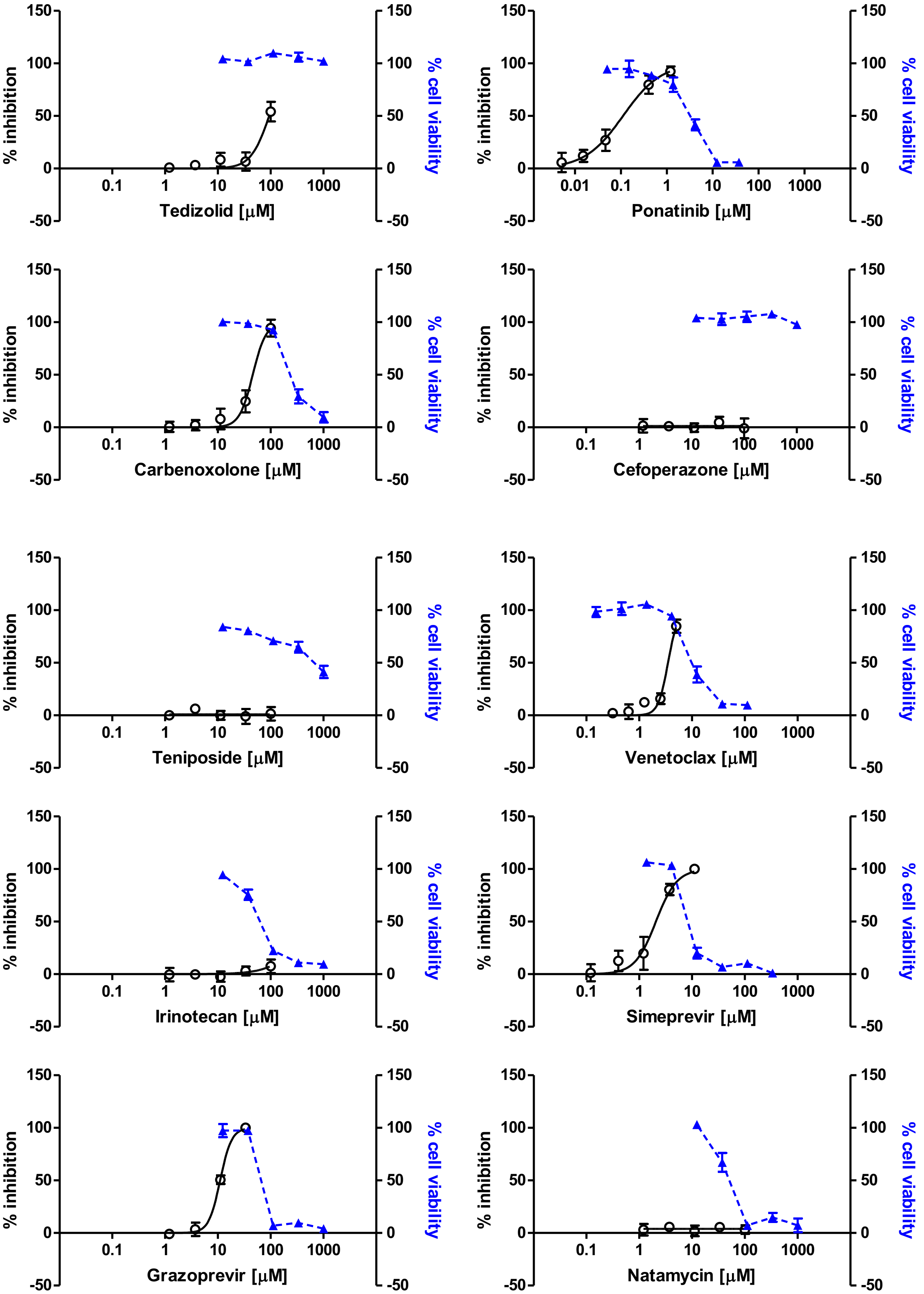
% inhibition
% cell viability
150
100
50
0
-50
0.1
1
10
100
1000
Tedizolid [μM]
0.01
0.1
1
10
100
1000
Ponatinib [μM]
Carbenoxolone [μM]
Cefoperazone [μM]
Teniposide [μM]
Venetoclax [μM]
Irinotecan [μM]
Simeprevir [μM]
Grazoprevir [μM]
Natamycin [μM]

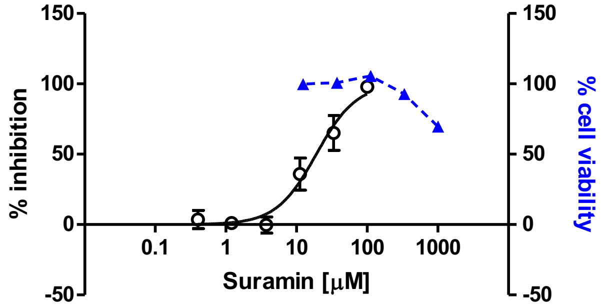


Figure S2: Antiviral efficacy of the selected compounds against HCoV-OC43. The antiviral activity of compounds was evaluated by means of focus reduction assay, infecting MRC-5 cells in presence of increasing concentration of compounds. Cell viability assays were performed in the same conditions as for antiviral assays, in absence of viral inoculum. The percentage infectivity inhibition (white dots) and the percentage of cell viability (blue triangles) were calculated by comparing treated and untreated wells. Error bars represent the standard deviation (SD) of 3 independent experiments.
